# Robust Segmentation of Cellular Ultrastructure on Sparsely Labeled 3D Electron Microscopy Images using Deep Learning

**DOI:** 10.1101/2021.05.27.446019

**Authors:** Archana Machireddy, Guillaume Thibault, Kevin G. Loftis, Kevin Stoltz, Cecilia E. Bueno, Hannah R. Smith, Jessica L. Riesterer, Joe W. Gray, Xubo Song

**Affiliations:** Computer Science and Electrical Engineering, Oregon Health and Science University, Portland, OR, 97239 USA; Department of Biomedical Engineering, Oregon Health & Science University, Portland, OR, 97239, USA; Multiscale Microscopy Core, Oregon Health & Science University, Portland, OR, 97239, USA; Department of Medical Informatics and Clinical Epidemiology at Oregon Health and Science University, Portland, OR, 97239 USA; Knight Cancer Institute, Oregon Health & Science University, Portland, OR, 97239, USA

**Author notes:** Further information should be directed to the Lead Contact, Xubo Song.

## Abstract

A deeper understanding of the cellular and subcellular organization of tumor cells and their interactions with the tumor microenvironment will shed light on how cancer evolves and guide effective therapy choices. Electron microscopy (EM) images can provide detailed view of the cellular ultrastructure and are being generated at an ever-increasing rate. However, the bottleneck in their analysis is the delineation of the cellular structures to enable interpretable rendering. We have mitigated this limitation by using deep learning, specifically, the ResUNet architecture, to segment cells and subcellular ultrastructure. Our initial prototype focuses on segmenting nuclei and nucleoli in 3D FIB-SEM images of tumor biopsies obtained from patients with metastatic breast and pancreatic cancers. Trained with sparse manual labels, our method results in accurate segmentation of nuclei and nucleoli with best Dice score of 0.99 and 0.98 respectively. This method can be extended to other cellular structures, enabling deeper analysis of inter- and intracellular state and interactions.

## Introduction

Recent advances in tumor biology have shown that the plethora of interactions between tumor cells and their microenvironment can significantly influence cancer behavior and/or treatment responses by forming barriers that hinder drug access (Tanaka and Kano, 2018), providing chemical and mechanical signals that alter the functional states of tumor and stromal cells (Hirata et al., 2017) and influencing immune cell activity (Galli et al., 2020). A deeper understanding of the underlying cellular and molecular mechanisms intrinsic to tumor cells, as well as extrinsic influences from immune cells, other stromal cells, and structural microenvironments, will shed light on how cancer evolves and develops resistance to therapy (Thakkar et al., 2020). Understanding of these dynamic interactions can be used to develop novel approaches to disrupt key intercellular and intracellular interactions and facilitate the design and development of efficient therapeutic strategies to fight cancer (Baghban et al., 2020).

Multiple imaging technologies, including multiplex immunohistochemistry (Kalra and Baker, 2017), cyclic immunofluorescence imaging (Lin et al., 2018) and electron microscopy (EM) (Riesterer et al., 2020), have been used to map cellular states and inter- and intracellular interactions. EM is an important component in this imaging suite, providing nanometer resolution views of intra and intercellular interactions (Johnson et al., 2020) that are not apparent in images generated using light microscopy. This complete picture of spatial relationships can reveal potential therapeutic targets that can be related back to the macroscale heterogeneity and microenvironment of the tissue. Focused ion beam scanning electron microscopy (FIB-SEM) is especially informative, generating 2D stacks of serial SEM images that provide 3D information on subcellular features in large tissue volumes (Giannuzzi, 2005). FIB-SEM imaging proceeds via serial steps of SEM imaging of a sample surface and FIB removal of a uniform thin layer of the tissue with size comparable to the spatial resolution in x-y plane, thereby revealing a new surface to be imaged (Figure 1). This process is fully automated and ensures that imaged data is equidimensional in all three axes, which significantly improves the accuracy of feature recognition within the dataset (Bushby et al., 2011).

**Figure 1.**
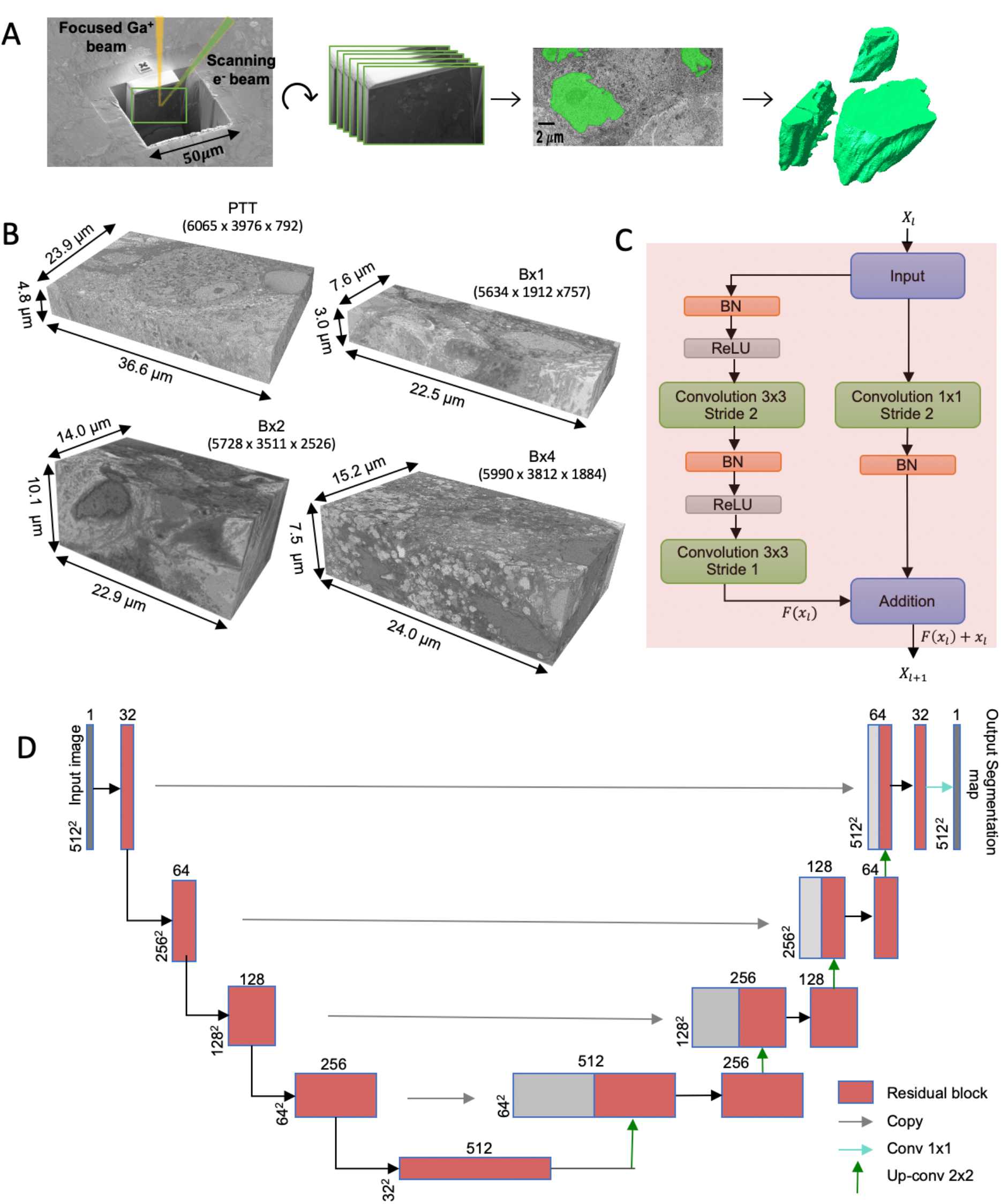
FIB-SEM-to-volume rendering workflow and ResUNet architecture. **(A**) Ga+ FIB-source sequentially slices a few nanometers from the sample to expose a fresh surface for subsequent imaging using the electron beam. An image stack was acquired and after image alignment and cropping, a small subset of the stack of images was segmented manually to generate a training set for the ResUNet. Once trained, the ResUNet was used to predict segmentation masks for the rest of the images in the stack. These predictions were used to create volume renderings for examination of 3D ultrastructural properties. (**B**) The four datasets and their sizes. The 3D FIB-SEM volume collected at 6 nm/voxel resolution from the biopsy sample (PTT) acquired from a patient with pancreatic ductal adenocarcinoma, and at 4 nm/voxel resolution from the biopsy samples Bx1, Bx2 and Bx4 acquired from a patient with metastatic breast ductal carcinoma, **(C)** Residual block used in ResUNet, BN stands for batch normalization and ReLU stands for rectified linear unit. *X_l_* and *X_l+l_* are the input and output features for the residual layer *l*, and *F* represents the residual function **(D)** ResUNet architecture, input size is written on the side of each box. The number of feature maps in each residual layer is written on top of each box.

SEM has been used in a number of fields including neuroanatomy, cardiology, diabetes, and kidney diseases. For instance, SEM is able to resolve the neuronal ultrastructure at nanometer resolution, providing new insights into the structural and functional aspects of brain (Zheng et al., 2018, Domínguez-Álvaro et al., 2019, Petrozzi et al., 2007, Gagyi et al., 2012). SEM has enabled the visualization of 3D structure of the collagen skeleton (Caulfield and Borg, 1979) and cardiomyocytes (Kanzaki et al., 2010), shedding light on the ultrastructure of the heart tissue and structural correlations to cardiac failure. Human pancreatic tissue has been analyzed with SEM to understand the pathophysiological mechanisms leading to type 1 diabetes (de Boer et al., 2020). EM has been widely used in the evaluation of kidney biopsies to diagnose hereditary nephritis, minimal change disease, and certain classes of lupus nephritis, which cannot be diagnosed based on light microscopy and immunofluorescence findings alone (Mokhtar and Jallalah, 2011, Shore and Moss, 2002).

SEM has also been used to examine cancer-derived cultured cells at nanometer resolution to gain insights, for instance, into cancer cell signaling networks (Pham and Ichikawa, 1992, López et al., 2017). The ultrastructure of inter- and intracellular interactions in 3D FIB-SEM images of human cancer biopsies have been explored by our group with emphasis on aspects of nanobiology that might affect therapeutic response (Johnson et al., 2020). Those studies reveal intricate interactions between tumor and stromal cells via micrometers long filopodia-like protrusions (FLP) suggesting the formation of the stomal barrier possibly driven by cytokine-receptor interactions that could be manipulated therapeutically to increase drug access and immune surveillance (Bazzichetto et at., 2019). The alignment of mitochondria along the length of an elongated cell and insinuated into nuclear folds increased their proximity which could alter aspects of DNA damage repair and reactive oxygen species signaling (Ogawa and Baserga, 2017; Saki and Prakash, 2017). A high abundance of lamellipodia and macropinosomes suggested nutrient scavenging via macropinocytosis as a possible tumor survival mechanism (Commisso et al., 2013; Hosios et al., 2016). A high prevalence of dense lysosomal vesicles has been suggested to play a role in reducing therapeutic efficacy in patients, as drug sequestration via lysosomotropism has been implicated as a mechanism of resistance to palbociclib and other CDK4/6 inhibitors (Villamil Giraldo et al., 2014). The presence of fenestrated nucleoli implied the possibility of increased ribosome-mediated protein production and the aberrant nucleoli could have contributed to aberrant protein synthesis and DNA repair (Orsolic et al., 2016). Deeper understanding of these inter- and intracellular ultrastructure and interactions revealed by EM imaging could help target the underlying signaling processes and potentially lead to the development of novel approaches to tumor therapy.

While 3D FIB-SEM images are being generated with ever increasing rate in ongoing clinical programs, the rate limiting step in their analysis is the delineation of the cellular structures to enable rendering of the images into interpretable forms. Currently, this is done by experts manually annotating images, and while effective, is extremely time consuming, tedious and dependent on the skill of the expert. The development of rapid, robust and automated machine learning methods to segment ultrastructural features is acutely needed for wide spread use of EM in large scale studies (Perez et al., 2014).

EM image segmentation in general is difficult due to similar intensity and texture variations across all parts of the image (Karabağ et al., 2020). The high density of ultrastructural features and the convoluted and intertwined nature of cancer and stromal cells makes it even more challenging (Johnson et al., 2020). The ultrastructure of tumor cells and the tumor microenvironment is very different from the widely studied neural and other normal cells (Baba and Câtoi 2007, Zink et al., 2004, Coman and Anderson., 1955). Therefore, segmentation methodologies designed for these cells cannot be directly applied for cancer cell ultrastructure segmentation (Shen et al., 2017, Linsley et al., 2018, Vergara et al., 2020, Müller et al., 2021). To overcome these difficulties, we propose a simple yet effective Residual U-Net (ResUNet) deep learning model for segmentation of cancer ultrastructure (Figure 1D). U-Net is the most prominent deep learning architecture used in medical image segmentation applications (Ronneberger et al., 2015, Machireddy et al., 2020). It consists of a contracting path that encodes the input image into feature representations and a symmetric expanding path which projects the discriminative features learned by the contracting path back to the spatial resolution of the input image to obtain a dense segmentation map. This unique architecture captures global image context while preserving spatial accuracy, thereby facilitating high-precision image segmentation (Ronneberger et al., 2015). However, as the depth of the U-Net increases, the gradient information propagated through the layers to update the network weights gets vanishingly small, thus preventing effective training of the network (Glorot and Bengio, 2010). The residual learning technique was proposed to solve this problem by adding a shortcut mapping to reuse data across subsequent layers such that each layer only fits a residual mapping. The gradients could easily backpropagate through the shortcut connection making optimization of deeper networks feasible and faster (He et al., 2016). The resulting ResUNet model architecture combines the strengths of feature concatenation from the contracting and expanding structure of U-Net and residual connections, facilitating propagation of local spatial information and ease the training of the neural network.

We apply ResUNet to 3D FIB-SEM images of tumor biopsies obtained from patients with metastatic breast cancer participating in the OHSU Serial Measurements of Molecular Architecture and Response to Therapy (SMMART) program or pancreatic cancer participating the Brenden-Colson Center for Pancreatic Care (Figure 1B). Additionally, the metastatic breast cancer imaging data shown here was generated as part of the NIH’s Human Tumor Atlas Network (HTAN) and is freely available. Leveraging the similarities of structures contained in a 3D EM image stack, ResUNet is trained on a small subset of manually labeled images evenly distributed in the 3D EM image stack to efficiently segment the rest of the images in the stack.

As a prototype, in this paper we apply the ResUnet network to segment nuclei and nucleoli from 3D FIB-SEM images as they are commonly used as cancer markers (Zink et al., 2004, Denais and Lammerding, 2014). It has been reported that changes in nucleus in terms of volume, surface, shape, density, ratio of nuclear volume to cytoplasm and ultrastructural characteristics such as invaginations and changes in chromatin observed in nuclei can aid in assessment of tumor malignancy (Baba and Câtoi 2007, Zink et al., 2004). The nucleolus, a distinct sub-cellular structure in the nucleus, is characterized in cancer cells by hypertrophy, numerical increase, micro-segregations, movement towards the membrane and formation of intranuclear canalicular systems between nuclear membrane and nucleolus (Baba and Câtoi 2007). Targeting these structural and functional changes in nucleoli is emerging to be a promising therapeutic approach for treatment of cancer (Lindström et al., 2018). To our knowledge, this is the first attempt in the field of cancer study to automate segmentation of cancer cell ultrastructure from 3D FIB-SEM images of human biopsies. While our prototyping effort is on nuclei and nucleoli, our method is extensible to other cellular structures, potentially allowing for complete segmentation of the 3D FIB-SEM image stack. This enables visualization and quantification of ultrastructural inter- and intra-cellular compositions and interactions, and makes it possible to not only identify potential nanoscale sites for targeted therapies, but also construct an ultrastructural atlas of developing cancer from normal to malignant.

## Results

### Model training setup

We evaluated the proposed ResUNet on 3D FIB-SEM images of three longitudinal tissue biopsy samples (Bx1, Bx2 and Bx4) acquired over three phases of cancer treatment from a single patient with metastatic ER+ breast ductal carcinoma and one biopsy sample (PTT) acquired from a patient with pancreatic ductal adenocarcinoma (PDAC). Extensive additional information about the three biopsies from the patient with metastatic breast cancer are available (Johnson et al., 2020). For each dataset, a ResUNet model was trained using a small subset of manually labeled images and the trained model was used to generate segmentation masks on the rest of the unlabeled images. We cropped the full EM image slices, typically of size around 6000×4000, into tiles of size 512×512, for computational tractability as well as for the purpose of generating enough training samples.

### ResUNet can accurately segment nuclei and nucleoli

The nuclei and nucleoli segmentation performance were measured using three metrics– the Dice score, precision and recall. To train the ResUNet, a sparse subset of training images (typically between 7 to 50 slices) was randomly selected from the 3D image stack and used to tune the network weight, by minimizing the difference between the ground truth segmentation and the network generated segmentation on these training slices. Once trained, the network was used to segment the rest of the slices in the stack that were not used for training, and a resulting performance metrics in terms of Dice score, precision and recall were calculated. This process is repeated ten times, and the mean performance metrics over the ten runs was reported. Figure 2B shows the performance metrics for PTT, Bx1 and Bx2 datasets (nuclei results in the top row, and nucleoli results in the bottom row). Depending on the datasets and the training set size, overall Dice scores of 0.95−0.99 and 0.70−0.98 were achieved for nuclei and nucleoli segmentation, respectively. In addition to the nucleoli boundary, the model accurately segmented the fenestrations within the nucleoli (Figure 3C). The performances for the Bx2 dataset were lower than the other two datasets, as the Bx2 image stack of 2526 images was deeper than for Bx1 and PTT which had images stacks of 757 and 792 images, respectively. As a result, the same number of training images constituted a much smaller percentage and thus less coverage of the content. Another cause for lower Dice score in nucleoli segmentation in the Bx2 dataset was the low quality of its ground truth labels. In fact, the segmentation produced by the ResUNet model was better and smoother than the ground truth (Figure 3C). However, benchmarking against a low-quality ground truth led to reduced performance values. Figure 3A displays the comparison between volumes rendered from manual annotations and those from predicted segmentation, with nucleoli nested inside the nuclei for PTT, Bx1 and Bx2 datasets. Figure 3B only displays the volume rendered from predicted segmentation for Bx4 as it does not have ground truth labels for the entire stack. Figure 3D displays the volume rendering of the fenestrations inside nucleolus in all four datasets. Figures 4 and 5 show 2D image slices overlaid with ground truth and predicted segmentation for PTT (Figure 4A), Bx1 (Figure 4B), Bx2 (Figure 5A) and Bx4 (Figure 5B). They illustrate the accuracy of model segmentation compared to the ground truth annotation. In Bx2 the model was able to correctly detect a nucleus that was not labeled in the ground truth as seen on the top left on slice 83 in Figure 5A.

**Figure 2.**
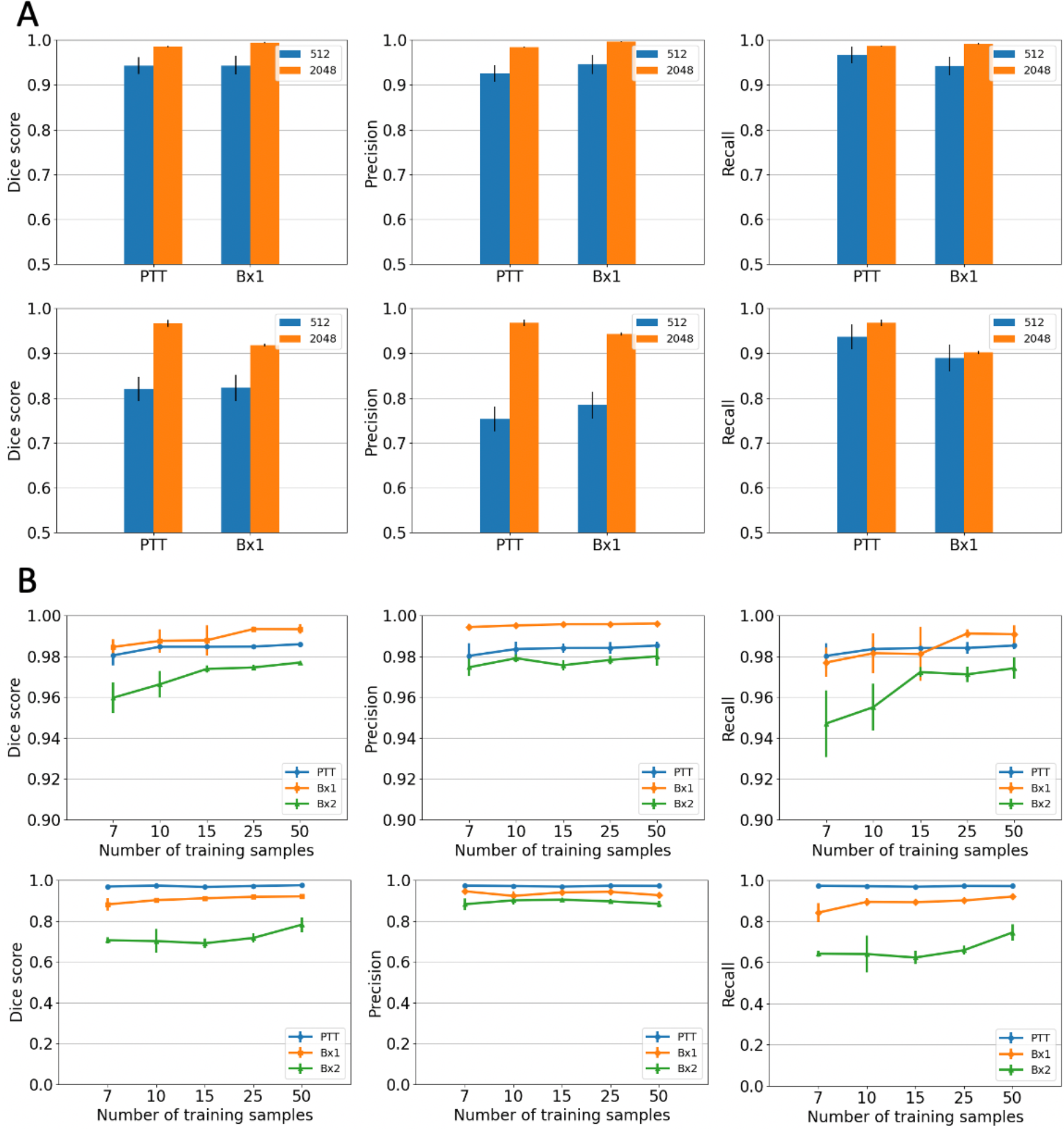
Nuclei and nucleoli segmentation performance (**A**) Effect of image tile size (context window). Segmentation performances for nuclei (top row) and nucleoli (bottom row) using different input tile sizes measured by Dice score (first column), precision (second column) and recall (third column) on the Bx1 and PTT datasets. The blue bar represents the results of training the network directly on the 512×512 sized tiles from the FIB-SEM images. The brown bar represents the results of training the network on image tiles of size 2048×2048 down-sampled to 512×512, which provide larger contextual information. Each error bar represents 10 separate experiments in which a new model was trained from scratch using the specified number of training images. (**B**) Effect of training set size. Segmentation performances for nuclei (top row) and nucleoli (bottom row) using different training set sizes, measured by Dice score (first column), precision (second column) and recall (third column) on PTT (blue), Bx1 (orange) and Bx2 (green) datasets. For each dataset, performance was evaluated over training set sizes of [7,10,15,25,50] in order to find the minimum number of images required to generate accurate segmentation. Each error bar represents the mean and standard deviation of results obtained from 10 separate experiments in which a new model was trained from scratch using the specified number of training images.

**Figure 3.**
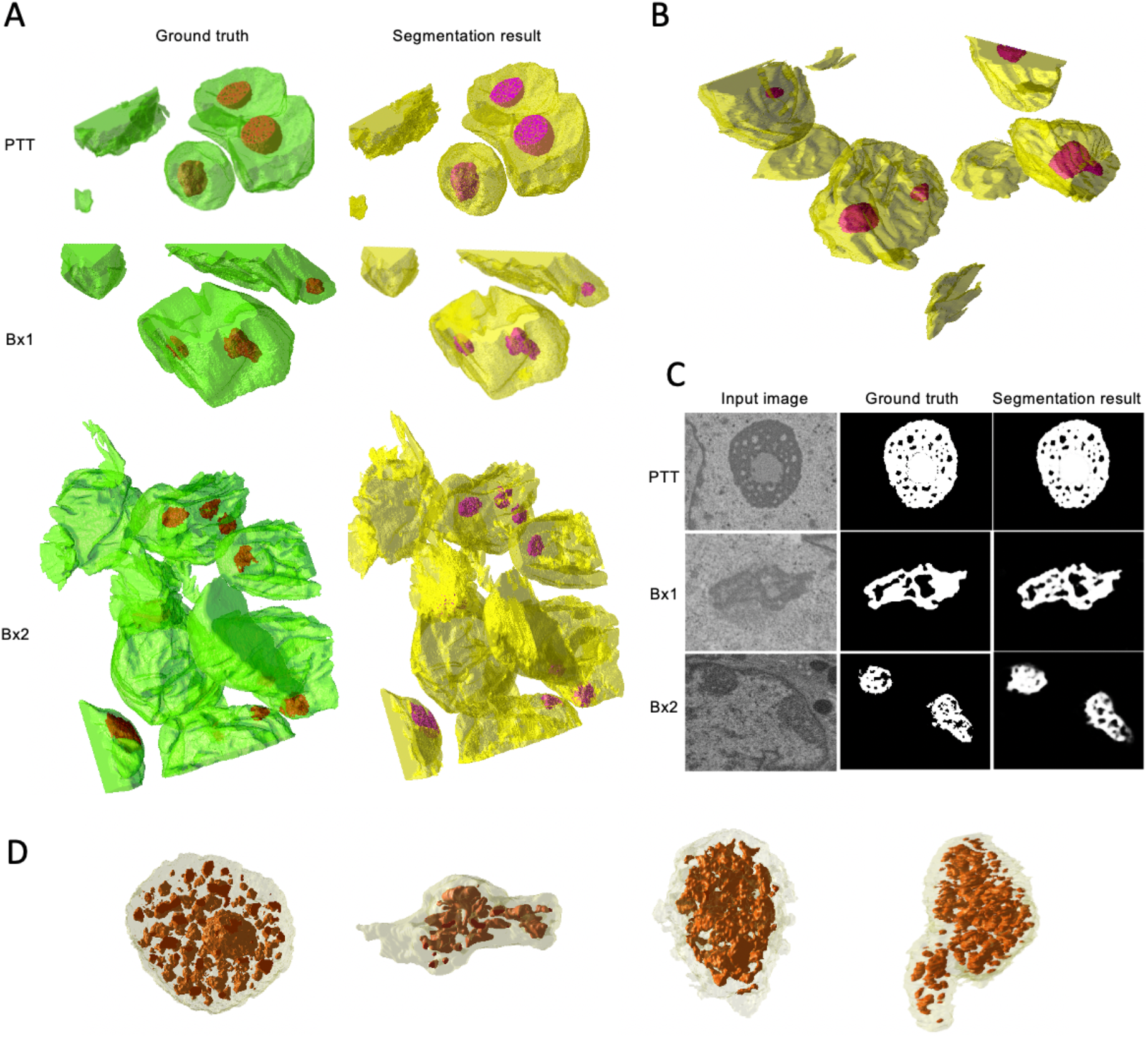
Nuclei and nucleoli volume renderings and nucleoli segmentation results (**A**) Volume renderings showing the 3D structure of ground truth masks and predicted segmentation masks for PTT, Bx1 and Bx2 datasets (**B**) Volume rendering showing the 3D structure of predicted segmentation masks for Bx4 dataset. (**C**) Representative qualitative results showing input images (first column), ground truth (second column) and predicted nucleoli (third column) for PTT (first row), Bx1 (second row) and Bx2 (third row) datasets. (**D**) Volume renderings of the fenestrations in nucleoli of PTT, Bx1, Bx2 and Bx4 datasets.

**Figure 4.**
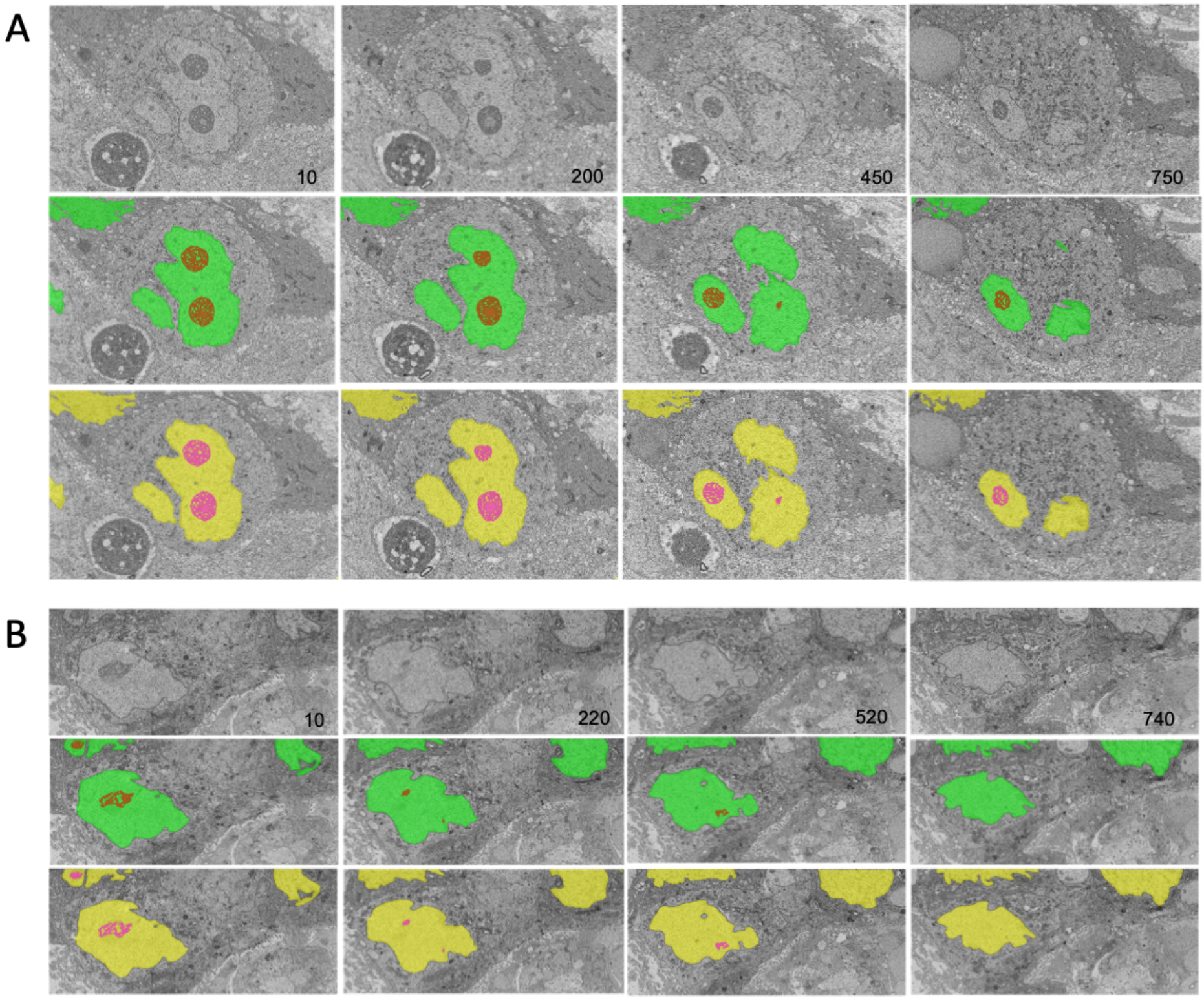
Nuclei and nucleoli segmentation results on PTT and Bx1 datasets. Representative qualitative results showing input images (top row) overlaid with nuclei (green) and nucleoli (red) ground truth masks (middle row) and predicted nuclei (yellow) and nucleoli (pink) segmentation masks (bottom row) for (**A**) PTT and (**B**) Bx1 datasets. Numbers in the lower right-hand corner of images indicate the slice position of the image in the full image stack.

**Figure 5.**
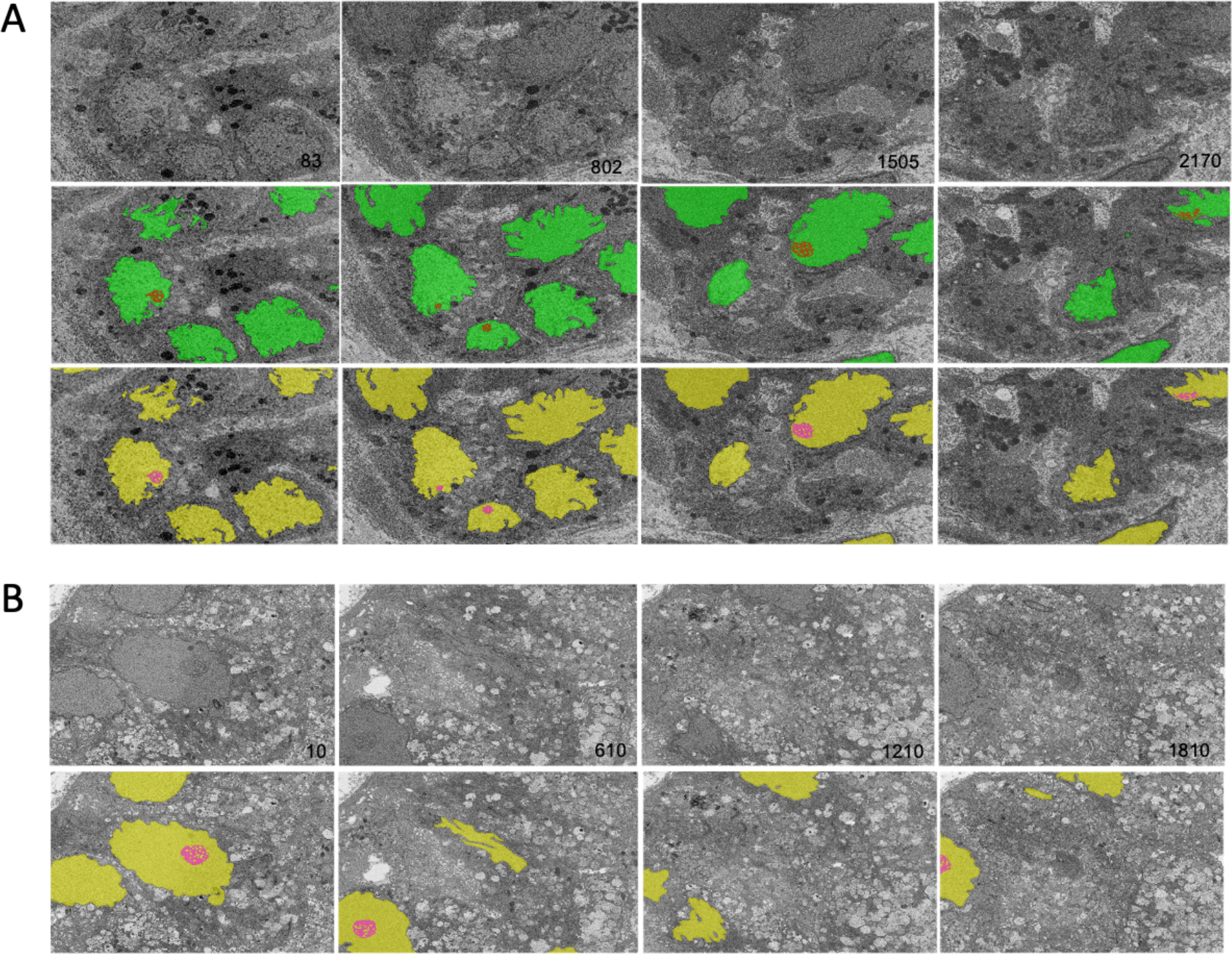
Nuclei and nucleoli segmentation results on Bx2 and Bx4 datasets. Representative qualitative results showing input images (top row) overlaid with nuclei (green) and nucleoli (red) ground truth masks (middle row) and predicted nuclei (yellow) and nucleoli (pink) segmentation masks (bottom row) for (**A**) Bx2 and (**B**) Bx4 datasets. Numbers in the lower right-hand corner of images indicate the slice position of the image in the full image stack.

### Larger context improves segmentation performances

The nuclei in cancer cells are relatively large. A 512×512 tile typically includes only a small section of the nucleus, and therefore, does not contain enough spatial context to cover the entire structure of the nucleus. We hypothesized that a larger crop size would provide better contextual information regarding the surroundings of nuclei and nucleoli and could lead to improved segmentation. In order to incorporate more global context while making it computationally feasible, we extracted tiles of size 2048×2048 and down sampled them to 512×512 and trained segmentation models on these down sampled tiles.

Providing larger context during training improved all three metrics in both nuclei (Figure 2A, top row) and nucleoli (Figure 2A, bottom row). The information provided by the larger context seems to outweigh any information loss due to down sampling. Further, using larger tile size enables faster processing of an entire EM image as fewer tiles would be required to reconstruct the final result.

### Small training set is adequate and larger training set brings further improvement

We experimented with varying the number of manually labeled full images used for training by selecting 7, 10, 15, 25 or 50 training images evenly distributed across the image stack. The models were trained on 2048×2048 sized tiles down sampled to 512×512 pixels. The model performed well with just 7 training images, with an overall dice score of 0.95−0.99 for nuclei (Figure 2B, top row), and a dice score of 0.70−0.98 for nucleoli (Figure 2B, bottom row), depending on the data sets. The performances continued to improve with more training images for all three metrics for both nuclei and nucleoli, reaching a dice score range of 0.97−0.99 for nuclei and 0.76−0.99 for nucleoli with 50 training images. Even with 50 training images, it was still a very small training set, representing only 2-5% of all the images in the stack. This illustrates that it is possible to train a reliable model with sparse manual labeling for 3D EM segmentation. The performances for the data set of Bx2 were lower than the other two data sets, for the reasons discussed above.

### Quantitative characterization of nuclei and nucleoli morphology and texture

Segmentation of nuclei and nucleoli allowed us to characterize biologically relevant features such as their morphology and texture. The morphological measures were designed to capture the size and shape, and include features such as solidity, sphericity, circular variance, percentage of volume fenestrated, and proximity of nucleolus to nuclear membrane. The texture features were designed to capture the spatial distribution of intensity patterns, and include features such as homogeneity, contrast, variance, correlation, and average pattern spectrum. The mathematical definitions of these features are provided in the Methods section.

Figure 6 shows the morphological features we extracted for nuclei (Figure 6A) and nucleoli (Figure 6B and 6C). The solidity feature, measuring the concavity of a surface, captures the nuclear envelope invaginations. The higher level of invaginations in Bx2 and Bx4 when compared to PTT and Bx1 (Figure 3A) is reflected by their lower solidity value (Figure 6A). Similarly, the relatively smooth envelope of nucleoli in PTT is reflected by its higher solidity value (Figure 6B). The solidity of the nucleolus was calculated by filling the holes in the volume to exclude the effect of the volume fenestrated by pores and quantify only the overall change in shape. The sphericity and circular variance features measure roundness of an object, and can be used to capture shape irregularities of nuclei and nucleoli, common characteristics of cancer cell. Additionally, for the nucleolus, we calculated the percentage of volume fenestrated by pores. These values are high for the PTT and Bx4 datasets as a result of the complex structure of pores within the nucleoli. Finally, varying levels of proximity of nucleoli to the nucleus membrane were also observed. This observation is consistent with published studies which suggest that nucleoli in cancer cells often move towards the nuclear membrane and form intranuclear canalicular systems between nuclear membrane and nucleolus (Baba and Câtoi 2007). Accordingly, the proximity of nucleolus to nuclear membrane feature captures the more centered positioning of nucleoli in the PPT dataset and the close proximity of nucleoli to nuclear membrane in the Bx2 dataset.

**Figure 6.**
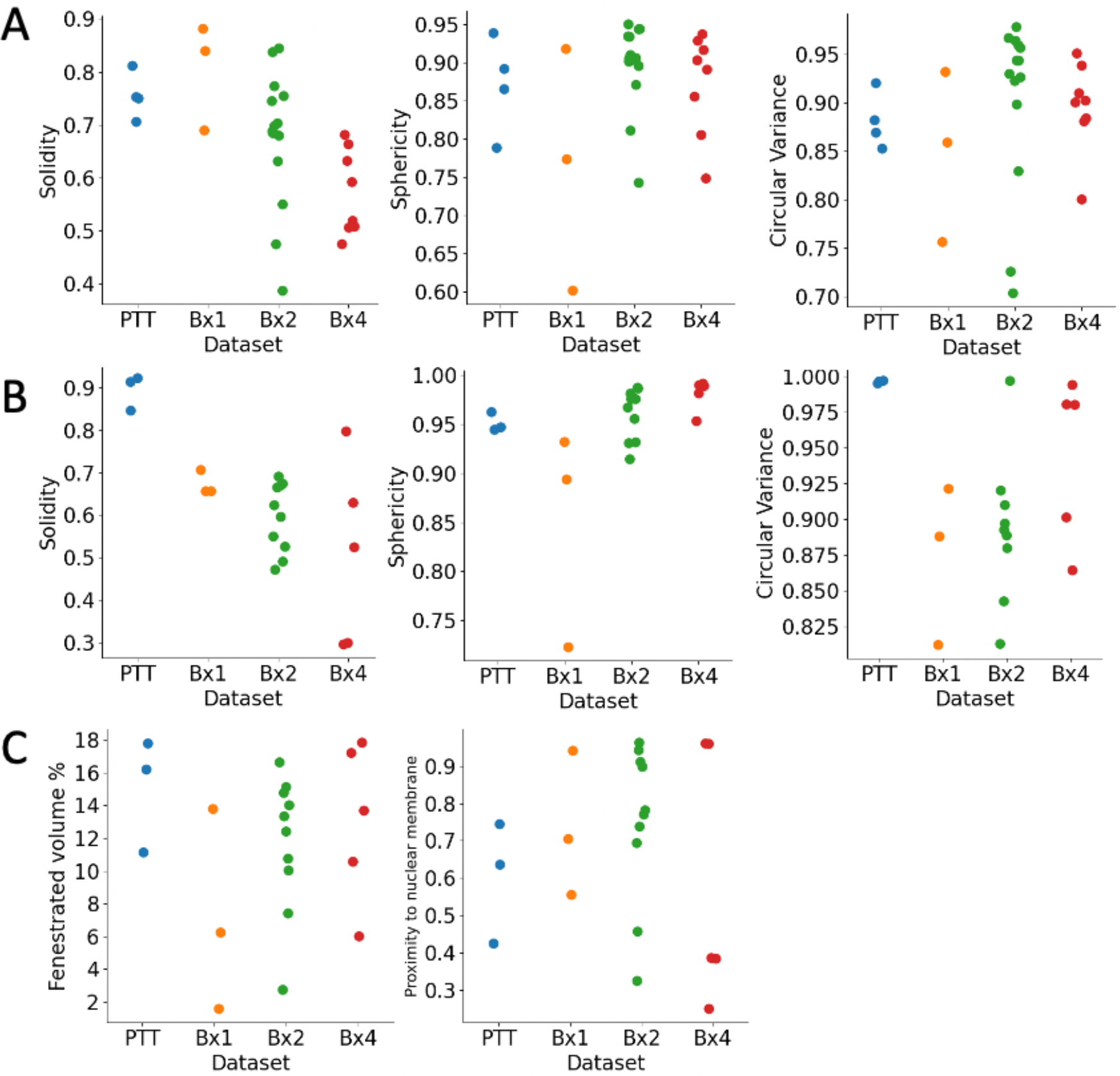
Morphological features extracted from nuclei and nucleoli. Solidity, sphericity and circular variance measures for (**A**) nuclei and (**B**) nucleoli in PTT, Bx1, Bx2 and Bx4 datasets. (**C**) Percentage of fenestrated volume in nucleoli and proximity of nucleoli to nuclear membrane for all datasets. Each dot represents the value of the feature for a nucleus or nucleolus.

Figure 7 shows the texture features we extracted for nuclei (Figure 7A) and nucleoli (Figure 7B). The texture features capture the spatial distribution of intensity patterns associated with chromatin and other internal structures. Texture characterization is well studied in the field of image processing. We use three classic methods, namely the grey level co-occurrence matrix (GLCM), the pattern spectrum, and the size zone matrix (SZM), to derive a set of texture features listed in Figure 7. GLCM captures the distribution of co-occurring gray-level intensities (Baraldi and Panniggiani, 1995). From GLCM, we extracted features such as homogeneity, correlation, variance and contrast for the four datasets (Figure 7A). The pattern spectrum feature characterizes the distribution of the sizes of various objects in an image (Thibault, et al., 2017). The SZM quantifies the size of homogenous grey level zones in an image (Thibault, et al., 2013), from which we extracted three features - zone sizes centroid, zone percentage and small zone high grey level emphasis. These features capture the sizes of groups of voxels of similar gray level intensities and can characterize the granularity of the chromatin structure in nuclei. These texture features are capable of capturing the differences between the images, and can be potentially used for downstream analysis.

**Figure 7.**
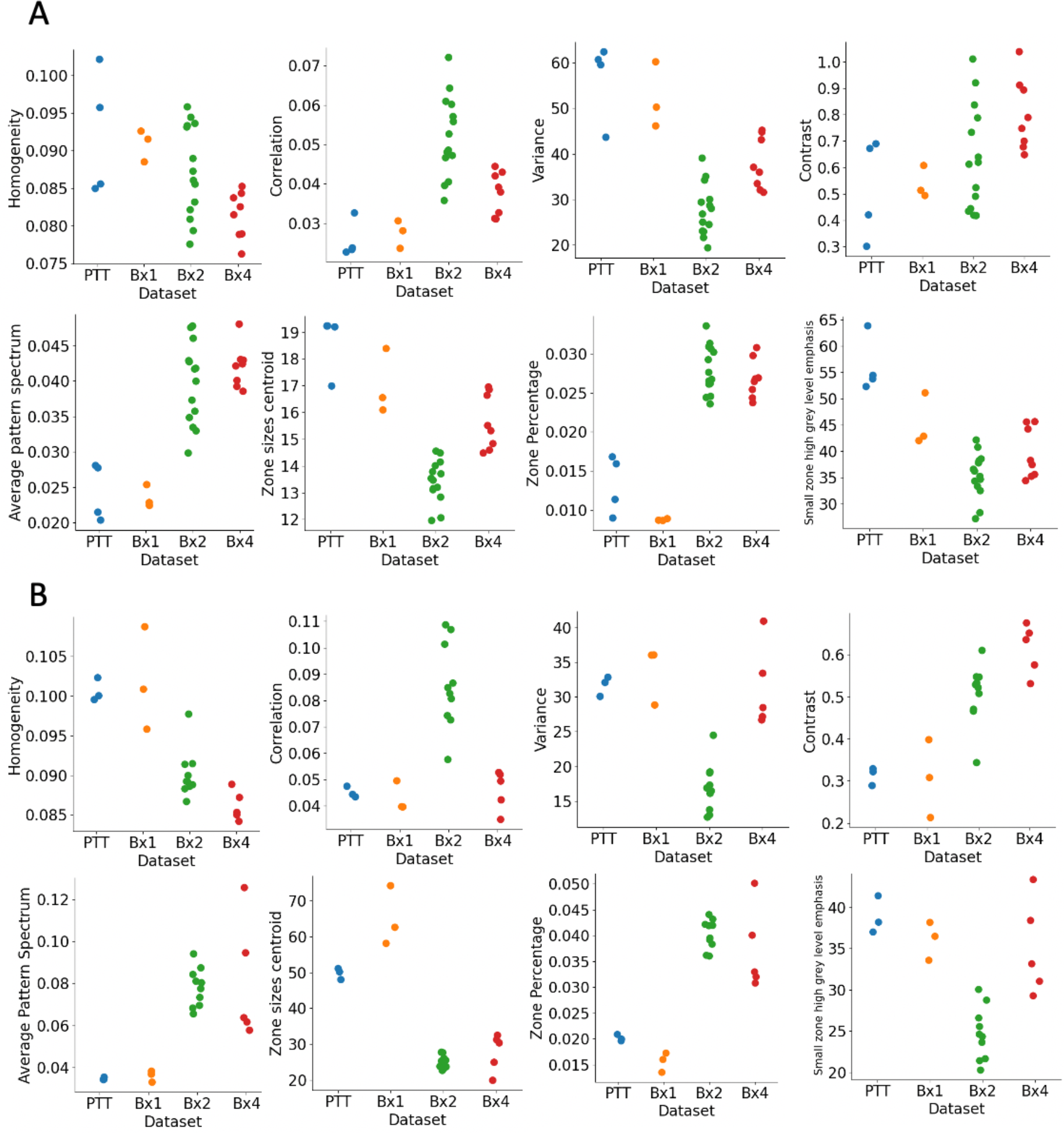
Texture features extracted from nuclei and nucleoli GLCM features (top row), pattern spectrum and SZM features (bottom row) for (**A**) nuclei and (**B**) nucleoli in PTT, Bx1, Bx2 and Bx4 datasets. Each dot represents the value of the feature for a nucleus or nucleolus.

## Discussion

We present here a deep-learning based approach for segmentation of cellular ultrastructure from 3D FIB-SM images, enabling rendering of the ultrastructure into interpretable forms. We used a ResUNet architecture to segment nuclei and nucleoli and evaluated the performance of the model on four human cancer biopsy samples from a clinical trial at OHSU supported by HTAN. The ResUNet architecture was trained with different sizes of training datasets to evaluate the number of manually labeled images required for accurate segmentation. It was observed that the number of manually labeled images required greatly depended on the variability of the structure across the dataset. When the structure was uniform across the dataset, roughly 1% of the dataset labeled was enough to train an efficient model. Even in the presence of large variations in the structure, 2% of dataset labeled seemed to be sufficient for good segmentation results. Also, using image tiles of size 2048×2048 down sampled to 512×512 improved segmentation results when compared to using 512×512 crops directly, as larger crops provided greater context which appeared to increase the segmentation performance. We demonstrated that ResUNet provided segmentation of nuclei with dice score 0.98, 0.99 and 0.97 and nucleoli with dice score 0.98, 0.92 and 0.74 for PTT, Bx1 and Bx2 datasets respectively.

Structures in 3D data collected via FIB-SEM exhibit high variability due to several factors, including the sample quality, tissue type, sample preparation techniques, microscope settings, and the imaging pixel resolution (Navlakha et al., 2013). In this study, we show that a small subset of the whole dataset contains enough information to capture most of the variability of a given structure with respect to the dataset, and demonstrate that a ResUNet trained with sparse labels is able to generate segmentation masks for the rest of the images in the stack. We also demonstrate the feasibility of morphology and texture quantification in nuclei and nucleoli. These quantitative features can be extracted efficiently, robustly, and reproducibly. While it is beyond the scope of this paper, we anticipate linking them to clinically relevant variables such as patient drug response in the future. This method can be extended to other cellular structures, enabling deeper analysis of inter- and intracellular state and interactions. The proposed segmentation of EM images fills the gap that has prevented modern EM imaging from being used routinely for research and clinical practices. It will enable interpretative rendering and provide quantitative image features to be associated with the observed therapeutic responses and answer diverse questions in biology.

## Acknowledgements

FIB-SEM data included in this manuscript were generated at the Multiscale Microscopy Core (MMC), an OHSU University Shared Resource, with technical support from the OHSU Center for Spatial Systems Biomedicine (OCSSB). The authors acknowledge the Knight Cancer Institute’s Precision Oncology SMMART Clinical Trials Program for the resources, samples, and data that supported this study. Specimen acquisition support from the SMMART clinical coordination team was invaluable. This manuscript was supported by the NCI Human Tumor Atlas Network (HTAN) Omic and Multidimensional Spatial (OMS) Atlas Center Grant (5U2CCA233280), Prospect Creek Foundation, the Brenden-Colson Center for Pancreatic Care, the NCI Cancer Systems Biology Measuring, Modeling, and Controlling Heterogeneity (M2CH) Center Grant (5U54CA209988), the OHSU Knight Cancer Institute NCI Cancer Center Support Grant (P30CA069533), and the OCS.

This study was approved by the Oregon Health & Science University Institutional Review Board (IRB #16113). Participant eligibility was determined by the enrolling physician and informed consent was obtained prior to all study protocol related procedures.

## Author Contributions

Conceptualization, J.W.G.; Methodology, A.M, G.T, X.S and J.W.G.; Software, A.M and G.T; Formal Analysis, A.M, X.S, G.T and J.W.G.; Investigation, A.M, X.S, J.L.R. and G.T; Resources, J.W.G; Data Curation, J.L.R, K.G.L, K.S., C.E.B and H.R.S; Writing – Original Draft, A.M and X.S; Writing – Review & Editing, A.M, X.S, G.T, J.L.R and J.W.G.; Visualization, A.M.; Supervision, X.S, G.T, J.L.R and J.W.G.; Project Administration, J.W.G; Funding Acquisition, J.W.G.

## Declaration of Interests

JWG has licensed technologies to Abbott Diagnostics; has ownership positions in Convergent Genomics, Health Technology Innovations, Zorro Bio and PDX Pharmaceuticals; serves as a paid consultant to New Leaf Ventures; has received research support from Thermo Fisher Scientific (formerly FEI), Zeiss, Miltenyi Biotech, Quantitative Imaging, Health Technology Innovations and Micron Technologies; and owns stock in Abbott Diagnostics, AbbVie, Alphabet, Amazon, Amgen, Apple, General Electric, Gilead, Intel, Microsoft, Nvidia, and Zimmer Biomet.

## STAR METHODS

### RESOURCE AVAILABILITY

#### Lead Contact

Further information should be directed to the Lead Contact, Xubo Song.

#### Materials Availability

This study did not generate new unique reagents.

#### Data and Code Availability

All primary datasets from clinical and exploratory analytics generated during this study are available through the HTAN Data Coordinating Center (https://humantumoratlas.org/) as patient HTA9_1 in the OMS Atlas.

### EXPERIMENTAL MODEL AND SUBJECT DETAILS

#### 3D Focused Ion Beam-Scanning Electron Microscopy Dataset Collection

Under an institutional review board approved observational study, three tissue biopsy samples (Bx1, Bx2 and Bx4) were acquired over three timepoints of cancer treatment from a patient with metastatic ER+ breast ductal carcinoma and one biopsy sample (PTT) was acquired from a patient with pancreatic ductal adenocarcinoma. Of note, Bx3 from the patient with breast cancer was not collected for EM analysis. All biopsies were acquired and analyzed under the OHSU IRB-approved protocol Molecular Mechanisms of Tumor Evolution and Resistance to Therapy (IRB#16113). HTAN sample IDs for Bx1, Bx2 and Bx4 are HTA9-1 bx1, HTAN9-1 bx2 and HTA9-5 (bx4) respectively. Informed written consent was obtained from the subjects. The samples were preserved in Karnovsky’s fixative (2.0% PFA, 2.5% Gluteraldehyde (Morris, 1965), post-fixed using an OsO_4_-TCH-OsO_4_ staining protocol, and embedded in Epon resin (Riesterer et al., 2020, López et al., 2017). Post-fixation staining binds heavy metals to lipid-rich membranes to provide contrast in EM imaging. Conductive coating with 8nm-thick carbon was necessary to achieve high-resolution, charge-free, high contrast, and low noise images. A FEI Helios NanoLab 660 DualBeam™ FIB-SEM was used to collect high resolution 3D volumes of the resin-embedded blocks. Targeted volumes were collected by using a Ga^+^ FIB-source to sequentially slice a few nanometers from the sample to expose a slightly fresh surface for subsequent imaging. The slicing/imaging cycle was automated using the FEI AutoSlice and View™ software extended package, while image collection during 3D data acquisition used the in-column detector (ICD). Metastatic breast cancer and primary pancreatic tissues were imaged with an isotropic resolution of 4 nm and 6 nm respectively.

#### Image preprocessing and ground truth generation

After data acquisition, images within the stack were translationally aligned in the XY-plane using an in-house stochastic version of TurboReg affine transformation (Thevenaz et al., 1998). The alignment step zero-padded the images in order to maintain a uniform size, which were subsequently cropped to yield the final 3D image volumes. The registration and edge cropping process yielded a final resolution of 6065×3976×792 for the PTT dataset, 5634×1912×757 for the Bx1 dataset, 5728×3511×2526 for Bx2 dataset and 5990×3812×1884 for Bx4 dataset. Intermittently, the brightness of a few images in the stack varied, increasing the complexity of the images and making segmentation more challenging. Histogram equalization was adopted to ensure consistency across the stack and reduce complexity of the images.

The nuclei and nucleoli were manually labeled using Microscopy Image Browser (Belevich et al., 2016) on all slices of PTT, Bx1 and Bx2 dataset and only 19 slices of Bx4 dataset. Manually labeling PTT, Bx1, and Bx2 datasets required roughly 50-80 hours each.

### METHOD DETAILS

#### Network Architecture

We combined residual blocks with a modified version of U-Net architecture. Similar to U-Net, the proposed network consists of an encoding path that extracts features, and a decoding path that up-samples the extracted features to obtain full-resolution segmentation, but uses residual blocks of convolutional layers as building units (Figure 1D). A residual block consists of two convolutional layers with a kernel size of 3×3 each preceded by batch normalization and rectified Linear Unit (ReLU) and a residual shortcut connection (Figure 1C).

The encoding path in the proposed network consists of four residual blocks. Instead of using pooling to down-sample the feature maps, a convolution with stride two is performed in the first layer of the residual block, which significantly reduces the computational load on the subsequent layers. The number of feature maps is doubled along each successive block in the encoding path to enable richer feature extraction. The encoding path is followed by a residual block which acts as a bridge between the encoding and decoding paths. Corresponding to the encoding path, the decoding path also consists of four residual blocks. The decoding path begins with the up-sampling of the feature maps found in the previous level of the decoding path, followed by a 2×2 convolution to half the number of feature maps. These feature maps are then concatenated with the feature maps at the same level in the encoding path through a skip connection. This concatenation step helps combine the deep, semantic, coarse-grained feature maps from decoder path with high resolution feature maps from encoder path enabling effective recovery of fine-grained details. These concatenated feature maps are then passed through the residual block. The output of the last residual block is passed through a 1×1 convolutional layer followed by sigmoid activation to provide the final segmentation mask.

#### Training data

The proposed ResUNet model was used to segment each 3D FIB-SEM stack by training on a small subset of manually labeled images and predicting on the remaining images. To estimate the number of manually labeled images required for efficient segmentation, the size of the training set was varied by selecting 7, 10, 15, 25 or 50 images evenly distributed across the dataset. The full resolution of the EM images cannot be analyzed directly by the network as it will be computationally infeasible and the resulting number of training samples for training will be too small. Therefore, image crop size of 512×512 was chosen in order to fit a meaningfully large number of images in a batch. We used a batch size of five in all our experiments, meaning that five image tiles were used to estimate the error gradient at each iteration of the network weight update. Using multiple training examples in a batch for error gradient estimation helps to make the training process more stable. We choose a batch size of five as it is the maximum number of tiles that could fit in the GPU memory in our case. The nuclei occupied 15-35% of pixels in each image and random selection of five 512×512 tiles per image ensured there was enough representation of nucleus in a batch. However, nucleoli are smaller and sparser, occupying less than 1% of the full image. As a result, if sampled randomly, a large number of tiles would need to be considered in order to form a batch with enough nucleoli representation, which would exceed the GPU memory. To ensure that the model encounters sufficient representation of nucleoli during training, the batch size was still maintained at five but the tiles were selected such that at least 4 out of the 5 randomly selected tiles contained no less than 100 pixels related to nucleolus. Horizontal and vertical flips were randomly applied as data augmentation steps.

#### Implementational details

The proposed network was implemented using Keras framework (Gulli and Pal, 2017) with TensorFlow (Abadi et al., 2016) as backend. The network was optimized by adaptive moment estimation (Adam) with 10^−4^ learning rate, exponential decay rates for moment estimates β1 = 0.9, β2 = 0.999, and epsilon = 10^−7^. It was trained to minimize the Dice loss function for 5,000 weight updates. The experiments were performed on a single NVIDIA Tesla P100 GPU.

#### Inference Methodology

As the size of an EM image is much larger than the 512×512 crop size processed by the network, each image was parsed into multiple overlapping tiles. Each tile was passed through the network to predict a probability for every pixel, and pixels with value greater than or equal to 0.5 were assigned to the foreground class. The resulting overlapping segmentation maps were blended by multiplying each map with a 2-dimensional tapered cosine window (a=0.5) and adding the result to reduce the edge artifact at tile borders.

### QUANTIFICATION AND STATISTICAL ANALYSIS

#### Evaluation metrics

The segmentation mask predictions were evaluated using three metrics: Dice coefficient, recall and precision. The Dice coefficient provides a measure of overlap between the detected output and the ground truth and is defined in Eq. 1. It ranges between 0 and 1, where Dice coefficient of 1 denotes perfect overlap and 0 denotes no overlap. Dice coefficient is defined as,

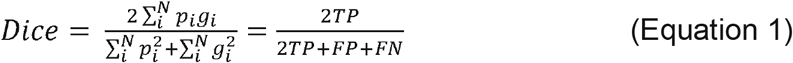

where *N* is the total number of pixels, *p* is the predicted pixel, *g* is the ground truth pixel, *TP* is true positives, *FP* is false positives and *FN* is false negatives. Recall is the fraction of the predicted pixels that actually belong to the foreground. If P is the predicted output, G the ground truth and ∩ denotes intersection, recall and precision are represented as,

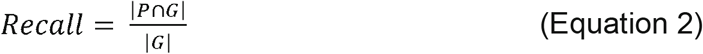

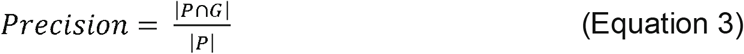

#### Morphological features

<u>Solidity</u> quantifies the concavities of a surface and is calculated as the ratio of the volume of the object to the volume of the convex hull (smallest encompassing convex polygon) of the object.

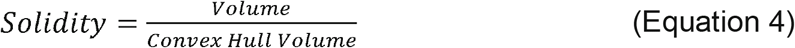

<u>Sphericity</u> is a measure of how close an object resembles a sphere. It is defined as the ratio of the surface area of a sphere with same volume as the object to the surface area of the object,

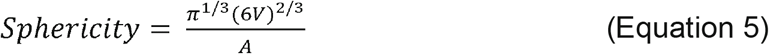

where *V* is the volume of the object and *A* is the surface area of the object (Wadell, 1935). Its value is 1 for a perfect sphere and decreases as the shape varies from a sphere.

<u>Circularity variance</u> provides a measure of the spread of radii across the volume. The lower the value, the tighter the clustering about a single mean. It is calculated as,

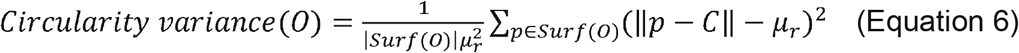

where *Surf* is the surface contour of the object *O*, *μ_r_* its mean radius and *C* its centroid.

<u>The percentage of fenestrated volume in the nucleoli</u> is calculated as the ratio of net volume to the filled in volume. The volume is filled in by performing a fill holes operation.

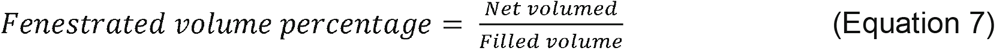

<u>Proximity of nucleolus to nuclear membrane</u> is calculated as the ratio of the distance between centroids of the nucleus and nucleolus (d_cc_) to the sum of d_cc_ and the distance between the centroid of nucleolus and the nearest point on the surface of the object (d_s_).

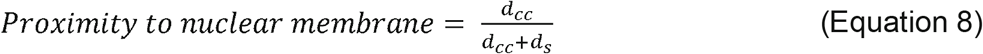

It takes a value of 0 when the nucleolus is at the center of the nucleus and increases as it moves towards the nuclear membrane.

### Texture Features

<u>Gray level co-occurrence matrix (GLCM)</u> examines the spatial relationship among pixels and defines how frequently a combination of pixel intensities occur for a given offset. A GLCM matrix was constructed by averaging the matrices obtained over 26 different offsets (Thibault et al., 2017). Four Haralick’s features, namely, homogeneity, correlation, variance and contrast, were computed from the GLCM matrix.

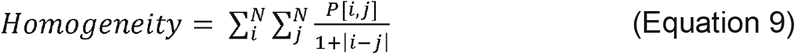

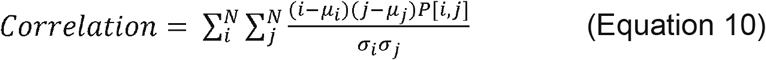

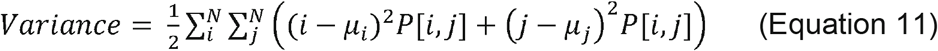

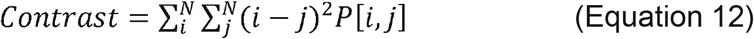

where *P[i,j]* is the *(i,j)*^th^ element of the GLCM matrix, *N* the number of gray levels and μ and σ denote the mean and standard deviation. <u>Homogeneity</u> measures the smoothness of the gray level distribution in the image. The homogeneity is 1 for a diagonal GLCM. <u>Correlation</u> measures the joint probability occurrence of certain pixel pairs and is high if the gray level of the pixel pairs is highly correlated. <u>Variance</u> gives a measure of dispersion of the gray level distribution and is large if gray levels are spread out. <u>Contrast</u> measures the local variations in the GLCM. Contrast value is low for smooth, soft textures and high for heterogeneous textures.

<u>Power spectrum (PS)</u> is a gray level rotation and intensity invariant texture descriptor based on granulometry. It is calculated by iteratively opening and closing an image with morphological operators and recording the resulting areas. It defines a probability distribution and the moments of this distribution are employed as signature patterns to characterize textures. It is calculated as,

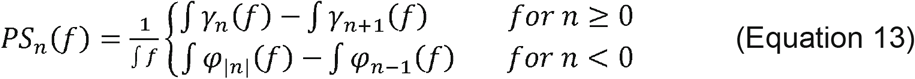

where *f* is the image, Γ= (*γ_n_)_n≥0_* and Φ = (*φ_n_)_n≥0_* are the morphological opening and closing operators and *n* is the size of the opening (Thibault, et al., 2017).

<u>Gray level size zone matrix (SZM)</u> provides a statistical representation of the clusters of gray levels in the image (Thibault et al., 2013). The value of the matrix *S(s_n_, g_m_)* is equal to the number of zones of size *s_n_* with gray level *g_m_* Three features, namely, zone percentage, small zone high gray level emphasis and centroid of zone sizes, are extracted from the SZM. <u>Zone percentage</u> measures the coarseness of the texture by taking the ratio of the number of zones (*N_z_*) to the number of voxels (*N*) in the image.

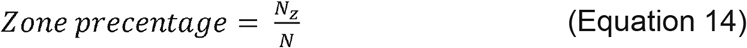

<u>The small zone high grey level emphasis</u> (SZHGE) measure highlights zone counts in the left quadrant of SZM where small zone sizes and high gray levels are located. The feature is defined as,

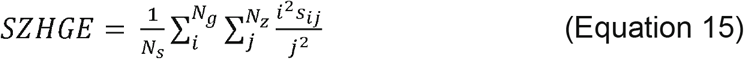

where *N_s_* is the total number of zones, *N_g_* is the number of discretized grey levels present in the image, *N_z_* the maximum zone size of any group of linked voxels, *s_ij_* is the number of zones with discretized grey level *i* and size *j*. The <u>centroid of zone sizes</u> is calculated as the centroid of SZM.

